# Data stochasticity and model parametrisation impact the performance of species distribution models: insights from a simulation study

**DOI:** 10.1101/2023.01.17.524386

**Authors:** Charlotte Lambert, Auriane Virgili

## Abstract

Species distribution models (SDM) are widely used to describe and explain how species relate to their environment, and predict their spatial distributions. As such, they are the cornerstone of most of spatial planning efforts worldwide. SDM can be implemented with wide array of data types (presence-only, presence-absence, count…), which can either be point- or areal-based, and use a wide array of environmental conditions as predictor variables. The choice of the sampling type as well as the resolution of environmental conditions to be used are recognized as of crucial importance, yet we lack any quantification of the effects these decisions may have on SDM reliability. In the present work, we fill this gap with an unprecedented simulation procedure. We simulated 100 possible distributions of two different virtual species in two different regions. Species distribution were modelled using either segment- or areal-based sampling and five different spatial resolutions of environmental conditions. The SDM performances were inspected by statistical metrics, model composition, shapes of relationships and prediction quality. We provided clear evidence of stochasticity in the modelling process (particularly in the shapes of relationships): two dataset from the same survey, species and region could yield different results. Sampling type had stronger effects than spatial resolution on the final model relevance. The effect of coarsening the resolution was directly related to the resistance of the spatial features to changes of scale: SDM failed to adequately identify spatial distributions when the spatial features targeted by the species were diluted by resolution coarsening. These results have important implications for the SDM community, backing up some commonly accepted choices, but also by highlighting some up-to-now unexpected features of SDM (stochasticity). As a whole, this work calls for carefully weighted decisions in implementing models, and for caution in interpreting results.

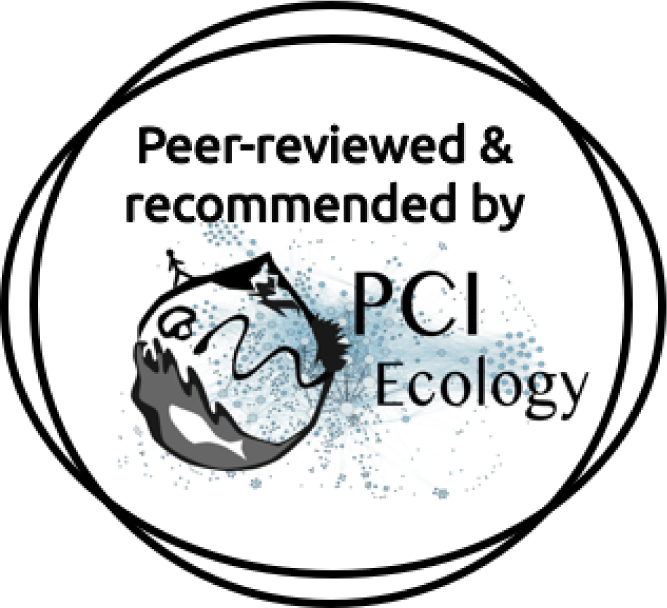

## 1 Introduction

Species distribution models (SDMs) are techniques widely used to describe and explain how species relate to their environment, often with the alongside goal of predicting their spatial distributions (Elith & Leathwick, 2009; Franklin, 2010). Thanks to the rapid growth of modelling techniques during the past decades, SDMs can today draw the best out of a wide array of data types, from presence-only data collected through citizen science to presence-absence data collected with standardised protocols (Guillera-Arroita et al., 2015). SDMs can be built on both direct (visual, acoustic, movement) or indirect (faeces, tracks) observations.

Henceforth, we have a good understanding of the statistical aspects underlying the various SDM methods available. We know how SDMs are sensitive to the number of observations underpinning a model (Hernandez et al., 2006; Virgili et al., 2018), how the temporal resolution of the environment conditions the success of a SDM in identifying the drivers of species distribution (Scales et al., 2016; Mannocci et al., 2017) and the various strengths and weaknesses associated with particular data types and particular methods. All these technical studies have permitted fine tuning high quality models providing reliable estimations of species distributions under both current and future environmental conditions. As such, SDMs outputs are today the cornerstone of most of spatial planning efforts worldwide, and most particularly in the marine realm (Franklin, 2013; Guisan et al., 2013; Marshall et al., 2014).

Yet, SDM accuracy rely on some technical choices for which we still lack a clear understanding of their impacts on the reliability and efficiency of the final model. The first choice relates to the spatial resolution of the environmental conditions the species distribution will be assessed against. Most of the time, this choice is constrained by technical aspects and data availability. Although the catalogue of environmental products is today wider than ever, with virtually most of physical and biological processes relevant to species distributions quantified and described at various scales throughout the Earth, trades-off must often be made between spatial and temporal resolutions, the finer temporal scale being most of the time associated with coarser grain size.

The choice of the spatio-temporal resolution of environmental conditions to be used must be motivated by the ultimate goal of the study and a good knowledge of the relevant ecological processes. All SDMs are based on the association of observation points with environmental variables, and the scale of these variables directly conditions the scale of the ecological processes identified. When the interest of a SDM lies in large scales, such that global description of species ranges, the coarser grain size (both spatially and temporally) of bioclimatic variables is often to be preferred (Scales et al., 2016; Mannocci et al., 2017; Manzoor et al., 2018). On the opposite, when the interest lies in describing the relationship of individuals to their direct habitat (be it through occurrence, presence probability or movement), finer resolutions of variables describing the small-scale features of habitats are more relevant (Scales et al., 2016; Manzoor et al., 2018). The literature provides evidence for poorer quality of model fit and overall degradation of model performances when coarsening the resolution of environmental variables (Guisan et al., 2007; Gottschalk et al., 2011; Connor et al., 2017), mostly due to the fact that coarsening results in homogenizing the various features composing a land or seascape, ultimately pruning information from the data. This, however, seems to be less acute for heterogeneous landscapes and specialist species, for which even degraded data contains sufficient signals for the model to find it (Hernandez et al., 2006; Lauzeral et al., 2013; Connor et al., 2017).

Most of this literature relates to terrestrial species with small home range sizes, or to plants, and is based on areal-based approaches (see Moudry et al., 2023, for a recent review). Although the issue of temporal resolution has been subject of several studies in the marine domain (Fernandez et al., 2017; Mannocci et al., 2017; Lambert et al., 2022), we still poorly understand the effects of spatial resolution on the performances of SDM for wide ranging mobile species. Large marine predators for example (cetaceans, elasmobranchs, seabirds, turtles, large teleosts) distribute over ranges covering thousands of kilometres square, often performing basin-scale migrations. For such species, SDMs are often built from aerial or ship-based standardised large-scale surveys following systematic transect designs, where any animals sighted around the transect are recorded (Lambert et al., 2019). Most of the time, the status and behaviour of sighted individuals are unknown. Given the extent of their range and their extreme mobility, individuals might be seen within a sub- or non-optimal ocean patch, while on their way between two places of interest (*i.e.*, two swarms of prey). This discrepancy substantially complicates matters and identifying the most relevant resolution can be particularly challenging: in that case, choosing too fine a resolution might result in individuals being associated with a certain environmental feature while they actually are targeting a more distant one.

Another crucial, yet understudied, aspect of SDM implementation is the choice of the sampling resolution. Standardised surveys can be either plot-, transect- or point-based and permit the collection of presence-absence data (Buckland et al., 2023). Transect-based surveys may be split subsequently into smaller-sized sampling units, generally of homogeneous detection conditions. The size of the splitting can be arbitrary, but must be informed based on the ecological processes of interest. For example, for aerial-based marine surveys, observation transects, which often exceed the hundreds of kilometres in size, can be split into 10, 5 or even 1 km long segments depending on the process of interest (see for example, among many others: Mannocci et al., 2014a; Lambert et al., 2022). For SDM purpose, the environmental conditions are generally retrieved to the segment centroids, so that the choice of the sampling resolution is related to the environmental products used and the two scales are chosen as being as similar as possible. Classical presence-absence or count-based analysis workflows with such data then collate sighting information on segment centroids, thereby ignoring the true position of the observation relative to the segment and effectively transforming transect data into point data.

However, point- and segment-based sampling can also be analysed as areal-based, where the number of observations and the amount of effort are summarized on spatial grids of arbitrary resolution, which is often that of the environmental variables (see for example Mannocci et al. (2014a,b) and Lambert et al. (2014) for a same dataset analysed both ways). The main advantage of this approach is to reconcile the spatial resolutions of both the predictor and the response variables (Moudry et al., 2023), thereby increasing the rate of observation per sampling unit (*i.e.*, reducing the number of zeros in the model), but we lose the information related to the fine-scale spatial patterns of the species.

The choice of the sampling resolution and type is recognized as crucial within the community, yet we lack any quantification of the actual effects these choices may have. Most of the time, the choice is made arbitrarily based upon the knowledge of the system at hand. In the present study, we aimed to fill this knowledge gap and to provide clues as to how these choices may affect the final ability of SDMs to successfully identify the relationship of a species to its environment, and how do these models fare in mimicking the true species distribution.

Given the impracticality of studying these aspects into natural environment and wide ranging species, we set up a simulation framework with two putative marine species within two regions characterized by different environmental forcings (Figure 1). For each species and region, we simulated 100 possible distributions of individuals based on species-specific environmental preferences. Virtual standardised strip-transect surveys were conducted over these simulations. The simulated survey data were subsequently formatted following either segment-based or areal-based (hereafter, raster-based) approaches, and each type was associated with five different spatial resolutions of environmental variables (0.083, 0.17, 0.25, 0.50 and 1.00°). We analysed each version of the dataset using the same analytical workflow, from model selection to prediction. Statistical performances, model composition, shapes of identified relationships and predicted species distributions were finally compared across models, as well as confronted to the true species distributions and to the true relationships to the environment as to assess the ability of models to successfully identify ecological patterns.

**Figure 1.**
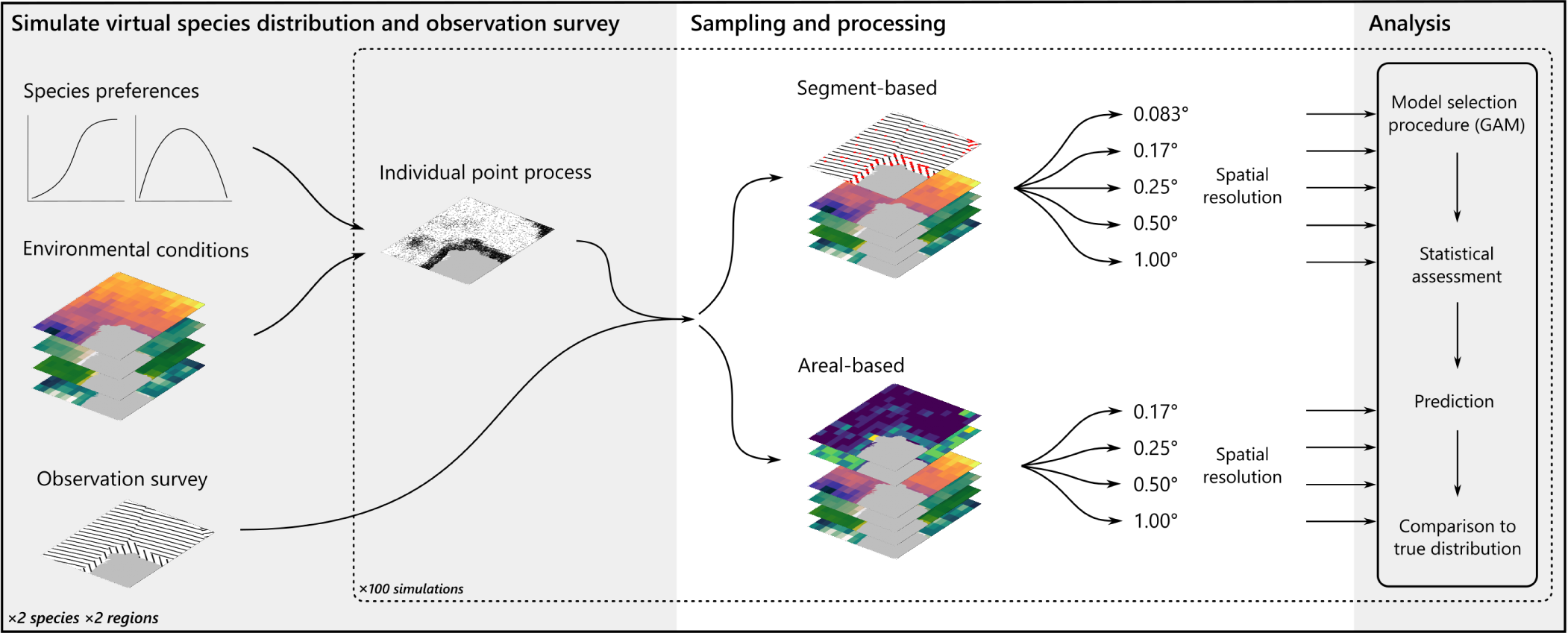
Conceptual workflow of the study. Four different species were built (two in two different study areas), and for each species the spatial distribution of individuals was generated 100 times. The observation survey was carried out over each simulated distribution, and observation data were formatted following segment-based and areal-based approaches. Each dataset was then associated with five different spatial resolutions of environmental variables, and all were analysed following the same analytical workflow.

## 2 Methods

All analyses were performed in R 4.0.4 (R Core Team, 2021).

### 2.1 Survey regions

We focused on the Eastern North Atlantic (ENA; more specifically, the western European shelf seas) and the Western Indian Ocean (WIO; more specifically, the Mozambique Channel).

The ENA region roughly corresponds to the Northeast Atlantic Shelves Province (Figure 2 and Supplementary Information S1 Figures 1–2; Longhurst, 2007), which is characterised by wide continental shelves and mega-tidal regimes. Oceanographic processes are dominated by tidal activity in the west, by freshwater inputs in the east, with wind mixing occurring throughout, generating an important spatio-temporal variability of local currents (Koutsikopoulos & Le Cann, 1996). The shelf edges are steep with strong slope currents flowing northwards, generating eddies in the southern Bay of Biscay (Pingree & Le Cann, 1992; Caballero et al., 2014). Tidal, freshwater and shelf edge fronts are thus dominating the marine region and sustaining its high productivity, even in winter.

**Figure 2.**
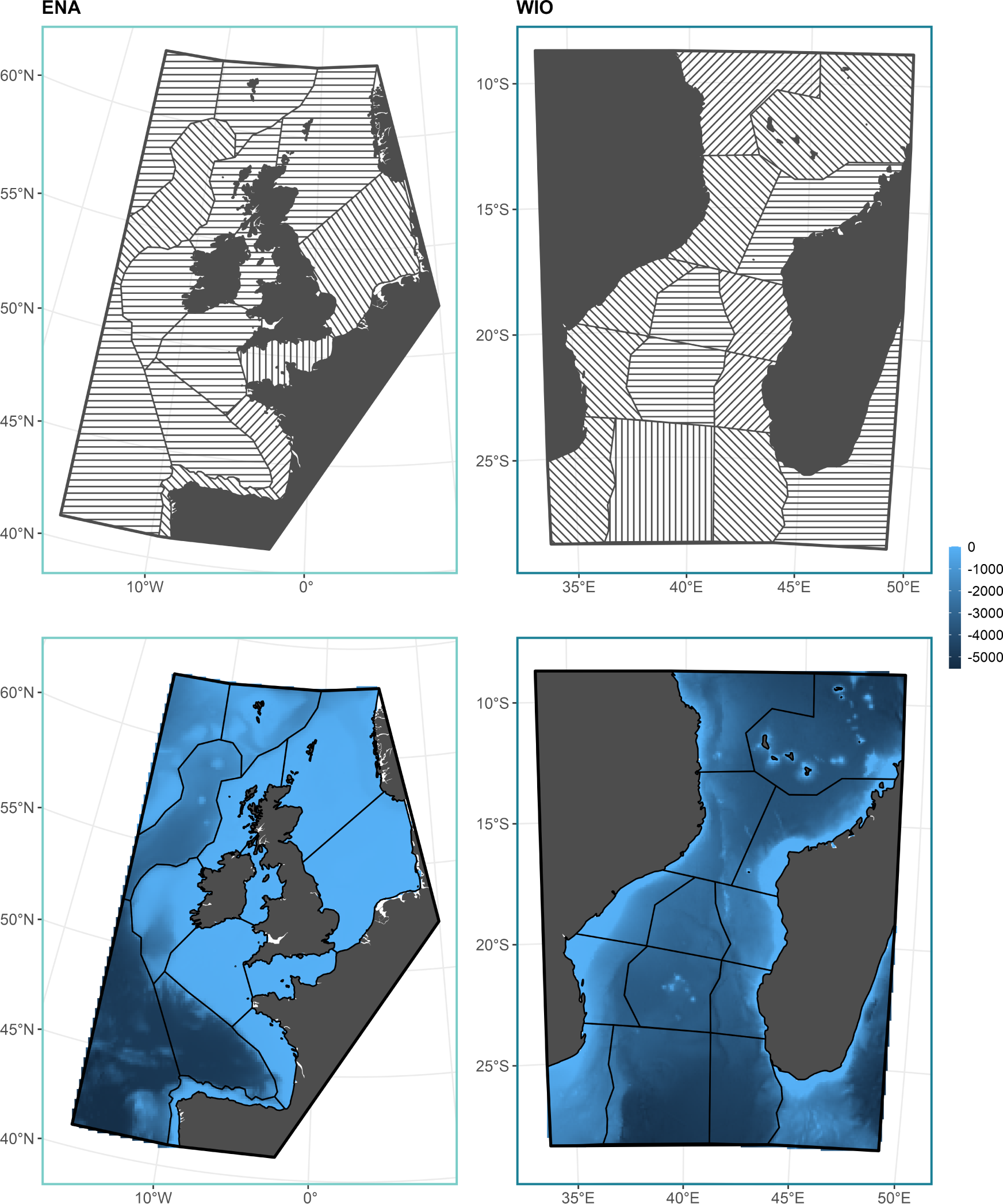
Study areas considered in the analysis: the Eastern North Atlantic (ENA, left) and Western Indian Ocean (WIO, right). Top panels display the survey strata and transects, the bottom panels display the bathymetric chart of both areas.

The WIO is an oceanic area (Figure 2 and Supplementary Information S1 Figures 2–3) mostly dominated by a strong mesoscale activity with a train of anticyclonic and cyclonic eddies flowing south through the Mozambique Channel from the anticyclonic cell around the Comoros (Schouten et al., 2003; de Ruijter et al., 2004). This meso-scale activity sustains most of the productivity in the area. The river plumes and relatively wide shelves off Mozambique and western Madagascar also induce important productivity along the coast, and probably participate in fuelling the biological enhancement associated with the eddies (Longhurst, 2007). In the north, the South Equatorial Current northern branch rounds Madagascar and interacts with the shallow banks of the Comoros to enhance productivity. In the south, the Mozambique eddies merge with those from eastern Madagascar to feed the Agulhas Current (Longhurst, 2007). Upwellings are also induced at the southern tip of Madagascar by the persistent anticyclonic eddy trapped there by topographic features.

**Figure 3.**
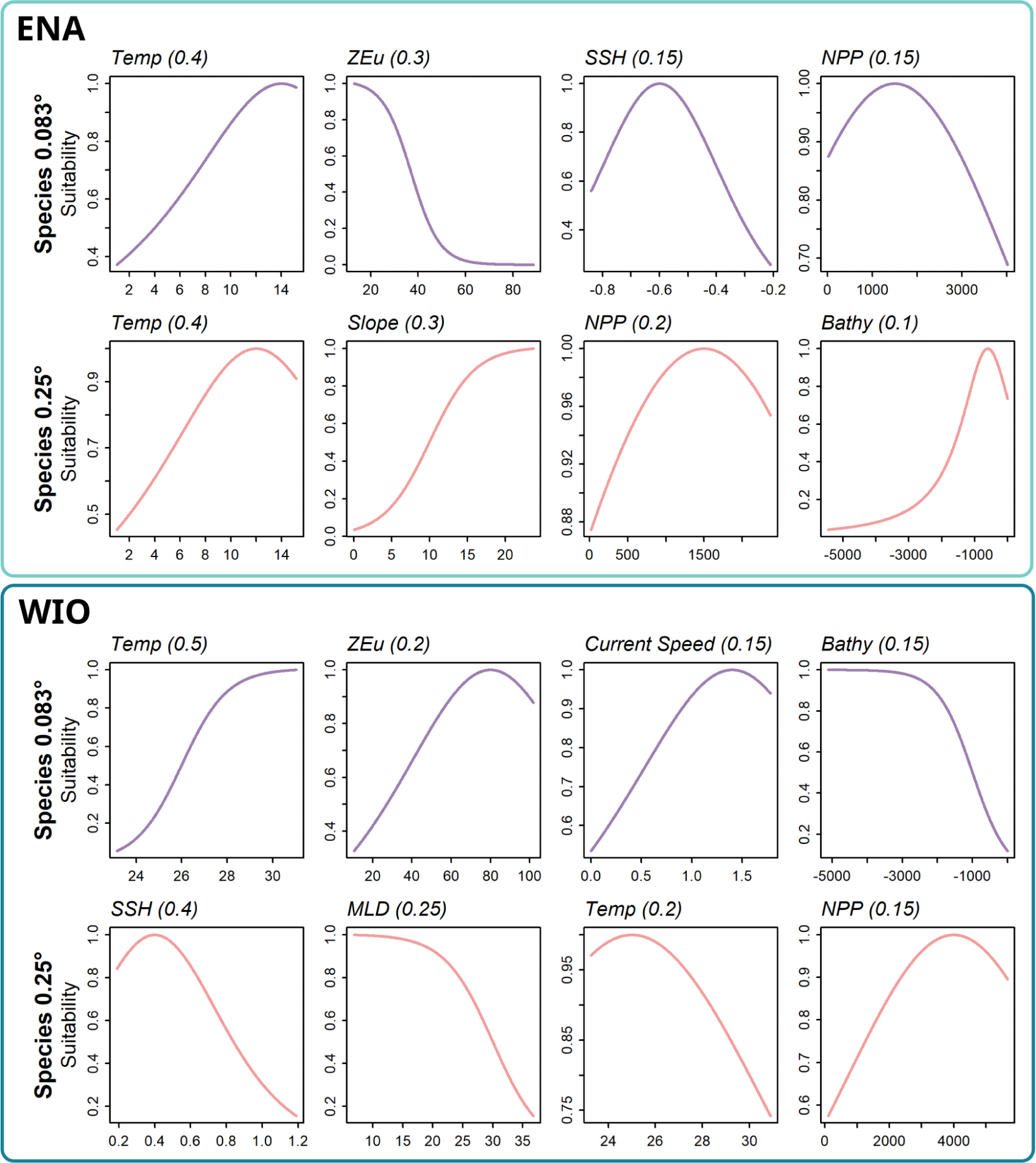
Response functions used to define the environmental suitability of each species in the ENA (top) and WIO (bottom). The weights of each variable in the species-specific additive equations are indicated in parenthesis. ENA: Eastern North Atlantic, WIO: Western Indian Ocean, Temp: sea surface temperature, ZEu: euphotic depth, SSH: sea surface height, NPP: net primary production, Bathy: bathymetry, MLD: mixed layer depth.

### 2.2 Oceanographic conditions

We extracted seven oceanographic variables from the Copernicus Marine Service (marine.copernicus.eu) in the study areas: sea surface temperature (Temp), sea surface height (SSH, mixed layer depth (MLD), euphotic depth (ZEu) and seawater velocity from the Global Ocean Physics Reanalysis (daily means). From the latter, we derived the current speed (CurrentSpeed) from the northward and eastward sea water velocities (computed as *sqrt*(*U* ^2^ + *V* ^2^), where *V* is the northward velocity and *U* the eastward velocity), and the Eddy Kinetic Energy (EKE; cm^2^.s^-2^; computed as 0.5*×*(*U* ^2^ +*V* ^2^)). The net primary production (NPP) was extracted from the Global Ocean Low-Mid Trophic Level Biomass Hindcast (SEAPODYM; daily mean). These variables were all extracted at the native 0.083° spatial resolution. For the purpose of this work, we made the choice of focusing on a single day, and arbitrarily choose the 2019, 1st February.

The GEBCO 2020 database was used to extract the bathymetry (hereafter, Bathy), from which we derived the Slope using the terrain function (from the raster package; Hijmans et al., 2014). These two variables were resampled at 0.083° to match the seven others.

We then aggregated all variables to coarser grain size by computing the mean of contiguous cells, aggregating them by factors 1.5, 3, 6 and 10 to obtain resolutions of 0.17, 0.25, 0.50 and 1.00°, respectively.

### 2.3 Simulating virtual species

We simulated two different species per region, whose distributions were driven solely by their fundamental niches. That is, their distributions were driven by environmental forcings alone, without any effects of biological factors like inter- and intra-specific interactions nor dispersal limitations. These environmental forcings were different for each species: one species distribution was driven by fine-scale processes (0.083° oceanographic conditions), the other by meso-scale processes (0.25° conditions). Response functions to four environmental conditions were built for each species, then combined to estimate environmental suitability maps further converted to occurrence probabilities, all using the virtualspecies package (Leroy et al., 2015).

All four species were built using a different set of variables and different response functions. In the ENA, the 0.083° species responded to Temp (with a Cauchy distribution), ZEu, SSH and NPP (with logistic distribution for all three). The 0.25° species responded to Temp, Bathy (with a Cauchy distribution for the two), NPP and Slope (with a logistic distribution for both). In the WIO, the 0.083° species responded to Temp, CurrentSpeed, ZEu and Bathy (all with a logistic distribution). The 0.25° species responded to SSH, MLD, NPP (with a logistic distribution) and Temp (with a Cauchy distribution).

We derived the environmental suitability from these response functions by additively combining them with species-specific coefficients for each variable (Figure 3). In the ENA, the suitability for the 0.083° was defined as 0.4 *× T emp* + 0.3*×ZEu*+0.15*×SSH* +0.15*×NPP*; for the 0.25°, it was defined as 0.4*×T emp*+0.3*×Slope*+0.2*×N PP* +0.1*×Bathy*. In the WIO, the suitability for the 0.083° was defined as 0.5 *× T emp* + 0.2 *× ZEu* + 0.15 *× CurrentSpeed* + 0.15 *× Bathy* (Figure 3); for the 0.25°, it was defined as 0.4 *× SSH* + 0.25 *× M LD* + 0.2 *× T emp* + 0.15 *× NPP*.

The environmental suitability was converted to occurrence probability through logistic conversion with a species prevalence of 10% (that is, the species occupy about 10% of the environment suitable to them; Figure 4). In the ENA, the 0.083° species mostly occurred within the Irish Sea, northern Channel and in southern Brittany, while the 0.25° was present throughout the shelf from the Irish Sea to the Iberian shelf, with the highest probability of occurrence found along the shelf edge in the Bay of Biscay (Figure 4). In the WIO, the 0.083° species mostly occurred along the edge of the cyclonic eddy in the Mozambique Channel, along the eastern coast of Madagascar and east of the Comoros while the 0.25° only occurred south of the Mozambique Channel (Figure 4).

**Figure 4.**
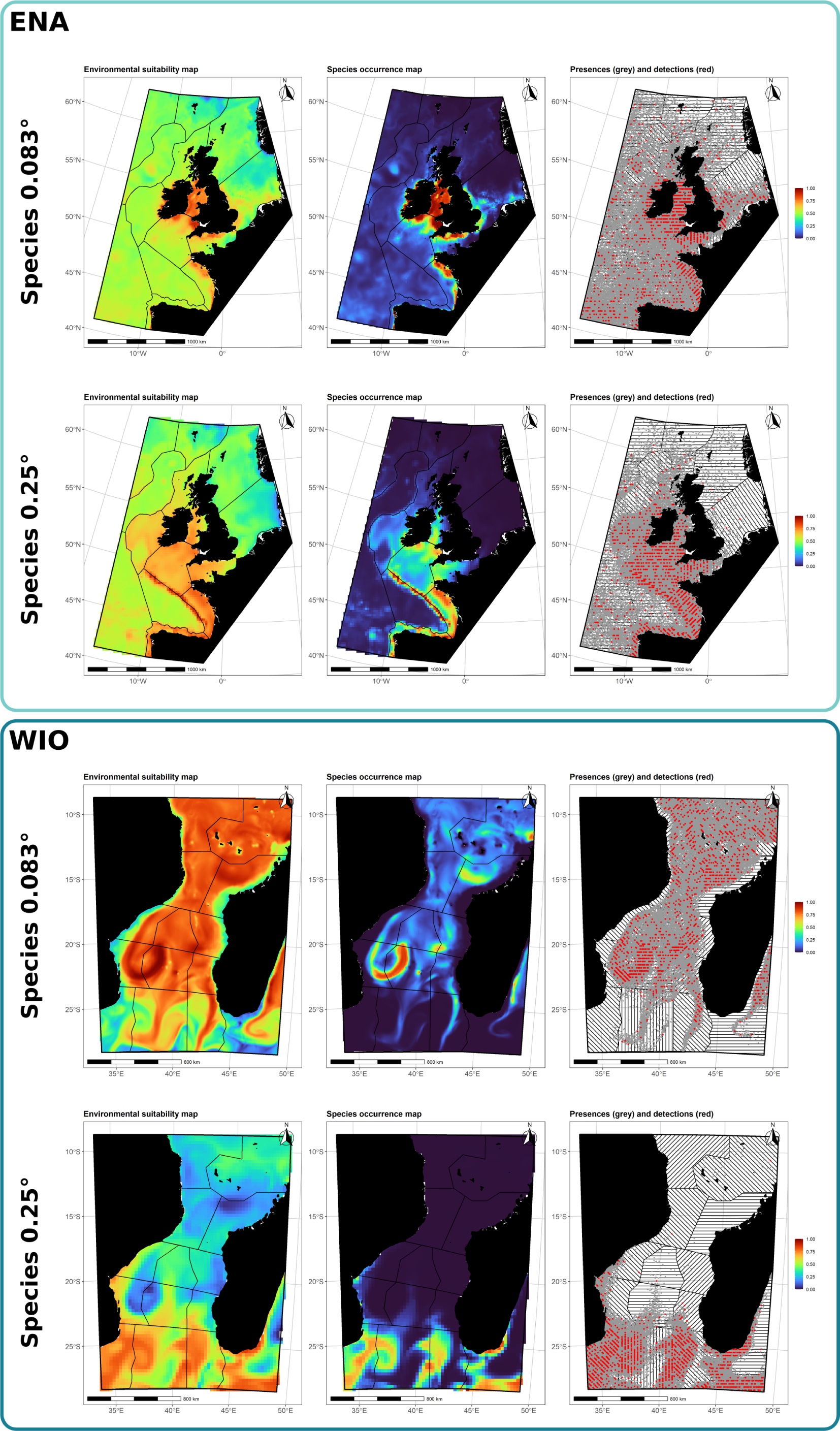
Environmental suitability derived from response functions (left), occurrence probability (center) and simulated distribution (one simulated survey; right) for each species in the ENA (top) and WIO (bottom). Grey points are simulated individuals, red points are individuals recorded during the survey and used for the subsequent analyses. ENA: Eastern North Atlantic, WIO: Western Indian Ocean.

We then generated 100 possible distributions for each species by building inhomogeneous Poisson point process (IPPP) using the species occurrence map as intensity parameter, and a population size of 100 000 individuals (using the spatstat package; Baddeley et al., 2015). This procedure permitted simulating the distribution of single individuals across the study area, with the number of individual per cell proportional to the occurrence probability (the actual position of individuals within each cell is random; Figure 4). We chose a population size for a common cetacean species. Using the same prevalence and population size for all four species avoided adding noise to our simulations, since the number of sightings directly affects SDM performance (Hernandez et al., 2006; Virgili et al., 2017, 2018).

### 2.4 Simulating virtual surveys

We simulated surveys following the Distance Sampling (DS) principles (Buckland et al., 2015), and the dssd package (Marshall, 2021). The overarching goal of DS surveys is to provide design-based density estimates; hence the survey design is a crucial step in their implementation. Given the size of our study areas, we used a multi-strata approach, with 14 strata in each region (each stratum was designed to be homogeneous in terms of bathymetry; Figure 2). The sampled transects followed a systematic design, assuming a maximum distance observation of 700 m, their position optimized to ensure a representative sample of the study regions and a uniform covering of each stratum (Figure 3). The transects were then divided into 10 km long segments.

We used the two survey designs to simulate virtual surveys over the 100 simulated distributions of the four species. We considered the surveys to be conducted over the course of a single day, the position of individuals to be static within each simulation and all individuals to be available for detection (that is, all individuals were considered to be at the sea surface). All individuals located within the 200 m around transects were considered as recorded by the survey, following the principle of strip-transect methodology (Figure 4; Buckland et al., 2015).

### 2.5 Sampling resolutions processing

To test the effect of sampling resolution on SDM performance, we formatted our simulated surveys into five segment-based resolutions and four raster-based resolutions. The segment-based resolutions were built by summarizing the number of individuals sighted by segment and associating segment centroids to the underlying values of oceanographic conditions with increasing grain sizes: 0.083°, 0.17°, 0.25°, 0.50°, 1.00° resolutions.

The raster-based resolutions were built by rasterising effort and observations of each simulation on the grids of oceanographic conditions: for each cell the total effort of segments whose centroid fell within was summed and the associated number of individuals sighted was summed for each species. This rasterisation was performed over the four coarser variable resolutions: 0.17°, 0.25°, 0.50°, 1.00°. The oceanographic conditions of the corresponding grid were then associated with each dataset. The 0.083° resolution being finer than that of survey segments (10 km), we did not construct raster-based dataset for this resolution.

### 2.6 Modelling

#### 2.6.1 Single variable approach

We adjusted single-variable Generalized Additive Models (GAM, with the mgcv package; Wood, 2006, 2011) to each variable, sampling resolution and simulation for the four virtual species. The models used the number of individuals sighted per sampling unit as response variable. They assumed a Tweedie distribution of residuals and used thin plate regression splines whose complexity was constrained to three inflection points maximum (*i.e.* four degrees of freedom maximum). This permits the GAM to adjust the complexity of the curve to the data, while avoiding overfitting. The number of individuals sighted per sampling unit was corrected by the area actually sampled within this unit by including this area as an offset in the model (2 *× segment length ×* 0.2 *m*). Predicted relationships between the number of individuals and each variable were stored for every simulation alongside summary statistics informing the quality of the model (explained deviance), the dispersion of residuals (RMSE) and the complexity of the fitted curves (estimated degrees of freedom).

These parameters were then compared across sampling resolutions and types (rasters *vs* segments), variables, species and regions. The differences in explained deviance, estimated degrees of freedom and RMSE between sampling types were computed for each simulation, sampling resolution, species and regions.

#### 2.6.2 Model selection and prediction

We implemented a regular model selection procedure for each simulation, sampling resolution, sampling type, species and region. The procedure tested every combination of up to four variables in the model, discarding all combinations of variables whose correlation was higher than 50%. Splines were bounded to four degrees of freedom maximum, and models were fitted with the REML method and the Tweedie distribution, including the sampled area as an offset (as above). The AIC and explained deviances were retrieved and stored for each tested model. Delta AIC and Akaike weights were then computed on models sorted by AIC (using the akaike.weights function from qpcR package; Spiess, 2018), and the first ranking model was selected as best model for that combination of simulation, sampling resolution and type, species and region. The variables selected in the best models were then compared to the original set of variables used to define the species distribution.

Species abundances were predicted from the selected best models. As to assess their reliability in reproducing the true species distribution, we used Pearson’s correlation coefficient (using the rcorr function from the Hmisc package; Harrell Jr et al., 2020) to compare each prediction to the distribution of true presences rasterised at the prediction resolution.

As to visually assess the predicted spatial pattern, we standardised every single prediction (by its maximum predicted density) before averaging them for each region, species, sampling resolution and type. We also summarised the true density of individuals on grids of the five resolutions (0.083°, 0.17°, 0.25°, 0.50°and 1.00°) for each simulation, and averaged them for visualisation purposes.

## 3 Results

### 3.1 Virtual surveys

The virtual surveys totalised 77 271 km of effort in the ENA, with an average over the 100 simulations of 1129 (sd 36.5) and 1107 individuals (sd 32.6) detected for the 0.083° and 0.25° species, respectively. In the WIO, a total of 79 959 km was sampled, for an average over the 100 simulations of 1475 (sd 38.1) and 1375 individuals (sd 37.4) detected for the 0.083° and 0.25° species, respectively.

When converted into 10 km-long segments, the number of sampling units for the segment-based models was 7819 in the ENA, and 8070 in the WIO. The rasterisation of sightings and efforts into target resolutions resulted in a sharp drop of the number of sampling units for raster-based models, alongside a sharp increase of the number of sightings per sampling unit (from an average of 16% to an average of 54% of sampling units with sightings, for all species and regions combined; Figure 5).

**Figure 5.**
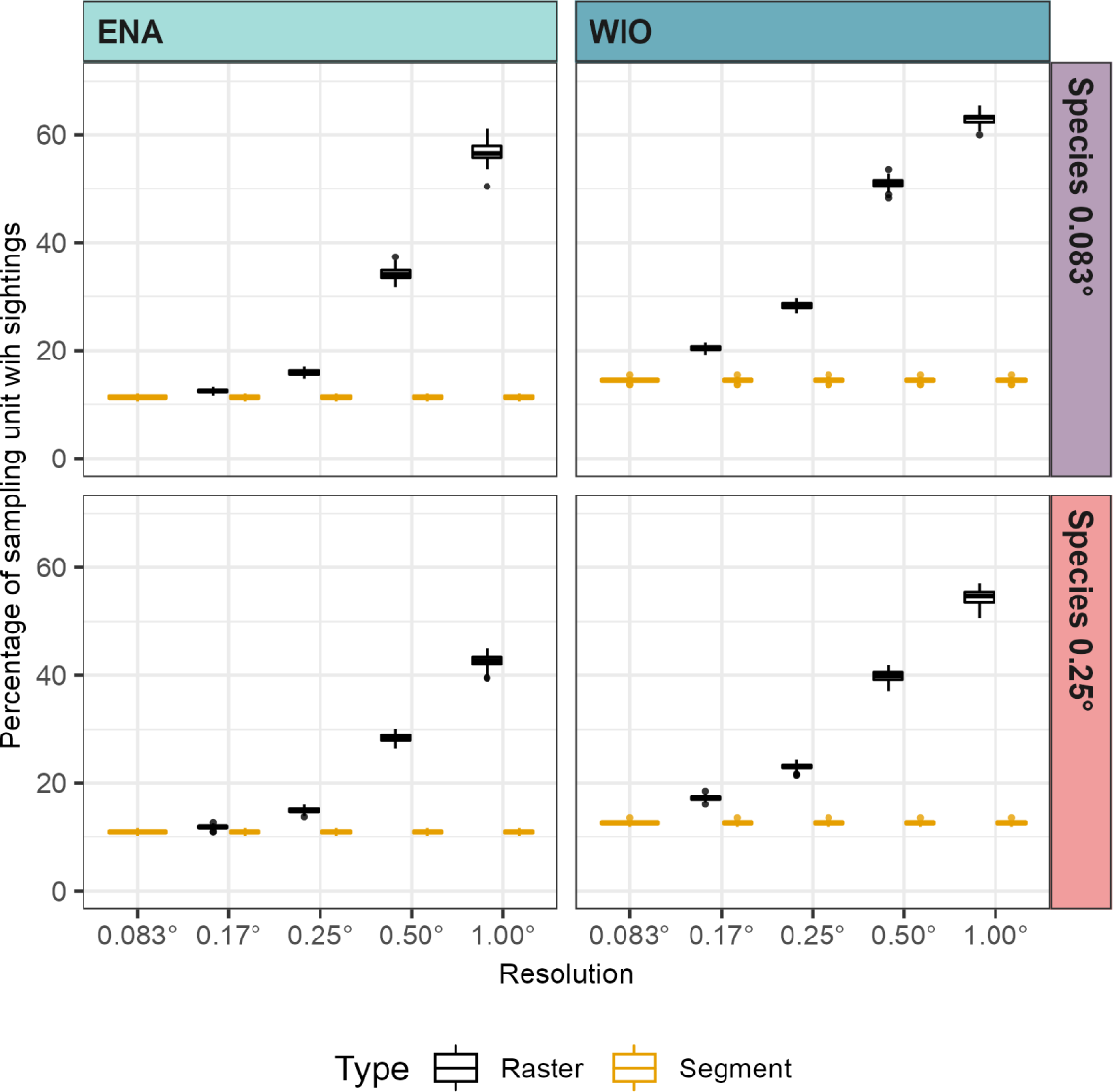
Proportion of sampling units with at least one individual detected for each sampling type and resolution, in both regions and species. ENA: Eastern North Atlantic, WIO: Western Indian Ocean.

When rasterised, the observed densities still displayed correctly the spatial patterns of occurrences for the ENA species (see Supplementary Information S1 Figure 4). However, part of the spatial pattern was distorted for the WIO 0.083° species at the largest resolutions. The spatial pattern of the WIO 0.25° species occurrence was mostly preserved as the resolution was enlarged, but the rasterised maps displayed medium densities in the northern and central parts of the study area despite the occurrence probably there being close to zero, actually reflecting the scarce presences occurring in the area.

### 3.2 Single-variable models

Segment-based and raster-based sampling yielded consistent results in terms of overall relative importance among variables (as of explained deviance, Figure 6A). In the ENA, NPP, ZEu, MLD, SSH and Bathy achieved the highest deviances for the 0.083° species; Temp, SSH and NPP followed by Slope, MLD and ZEu for the 0.025° species. In the WIO, Temp, NPP, CurrentSpeed, EKE, Bathy and ZEu had the highest deviances for the 0.083° species; Temp, SSH, then NPP and ZEu for the 0.25° species. The deviances were systematically larger for the raster-based sampling, but the deviances were also largely more variable compared to segment-based sampling (wider boxplots).

**Figure 6.**
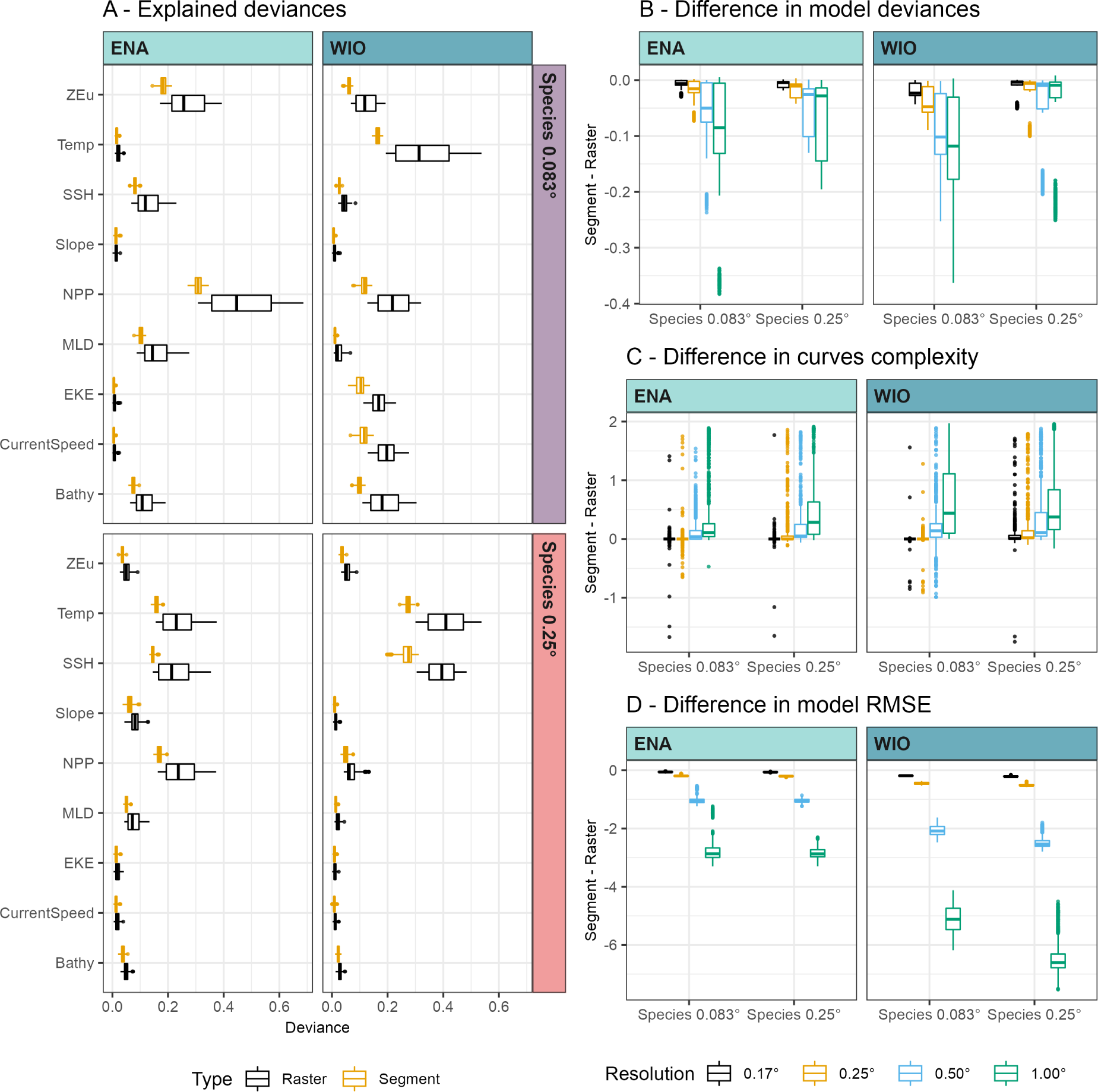
A - Distribution of explained deviances for single-variable models fitted to raster-based (black) and segment-based sampling (gold), for the two species and regions, whatever the sampling resolution (boxplots are constructed with each simulation occurring five times, one per resolution). B - Difference in explained deviances, C - curves complexity (estimated degrees of freedom) and D - RMSE between segment-based and raster-based sampling for each resolution and region. The differences are computed by simulations: one point behind the boxplots is the difference for a single simulation and a single variable between the dataset processed as segment-based *vs* raster-based (each simulation occur 9 × 2 times, one per variable and species). ENA: Eastern North Atlantic, WIO: Western Indian Ocean, ZEu: euphotic depth, Temp: sea surface temperature, SSH: sea surface height, NPP: net primary production, MLD: mixed layer depth, EKE: eddy kinetic energy, Bathy: bathymetry.

Computing the difference in deviance between the two sampling types for a single simulation and a single variable highlighted that the difference increased with increasing sampling resolutions, for all but the 0.25° species in the WIO (Figure 6B). The differences reached up to a 38% higher deviance for raster-based sampling compared to segment-based sampling with the 1.0° resolutions.

These higher deviances obtained with raster-based sampling and larger resolutions were probably linked to the higher sighting-to-effort ratio (Figure 5), and to a simplification of the fitted curves (Figure 6C): the difference of estimated degrees of freedom highlighted large and positive differences for larger resolutions (0.25 and 1.0°), whatever the species and regions, with an increasing proportion of linear curves for raster-based sampling at these resolutions. On the opposite, raster-based and segment-based sampling yielded similar curve estimations for the smaller sampling resolutions. Interestingly, the curve complexity was quite variable across simulations.

Accordingly, the RMSE was always larger for raster-based sampling, in particular for the 0.25 and 1.0° resolutions, indicating larger residuals and a poorer fit for raster-based than segment-based sampling at these resolutions.

### 3.3 Model selection

#### 3.3.1 Overview

The best models yielded quite variable results across simulations and within specific resolutions and types (Figure 7). Most of models were composed of two to four covariates for segment-based models, with some simulations selecting one variable only (0.50° and 1.00 ° resolutions for the 0.083° species in the ENA). Raster-based models were more variable in their composition across sampling resolutions, with several simulations selecting the null model as best one: 0.25° resolution for the 0.083° species in the ENA (53% of simulations), and the 0.17° resolution for the 0.083° species in the WIO (82% of simulations).

**Figure 7.**
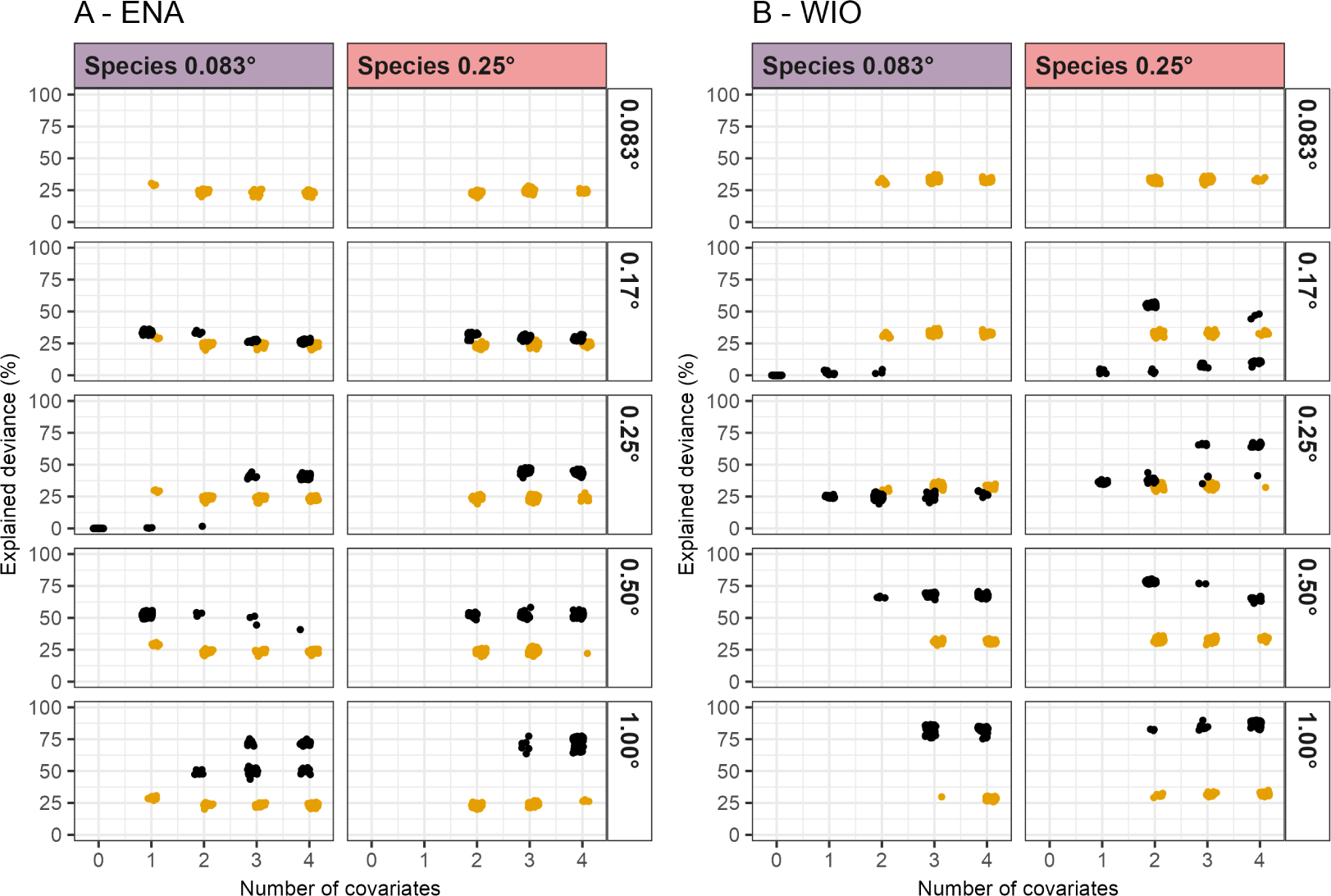
Relationships between explained deviances and numbers of covariates selected in best models fitted to raster-based (black) and segment-based sampling (gold), at the different resolutions for the two species and regions. ENA: Eastern North Atlantic, WIO: Western Indian Ocean.

The levels of explained deviances remained medium for most species and resolution (Figure 7; Supplementary Information S1 Figure 5), but with a large variability (ranging from 0 to 89%) mostly due to differences of performance across sampling types. Segment-based models explained deviances were quite stable across sampling resolutions, and did not vary according to the number of predictors in the model, suggesting that one or two main covariates sustained most of the explained deviance.

The raster-based explained deviances however showed wide variability across resolutions (Figure 7). Overall, the deviances tended to be higher for larger resolutions. For some species and resolutions, raster-based models had higher deviances than segment-based models (0.50 and 1.00° for both species in both ENA and WIO). In other cases, the selected raster-based models resulted in null explained deviances. This was mostly the case for simulations where the null model was selected, but also for some models selecting up to four predictors: 0.25° for the 0.083° species in the ENA; 0.17° for both species in the WIO. In the latter case, simulations with similar numbers of selected covariates were separated into two bulks of explained deviances, suggesting a variable model composition. Such bimodal distribution of simulations explained deviances within a single resolution also occurred for models reaching high deviances with large number of covariates (1.00° resolution for the 0.083° species in the ENA and the 0.25° resolution for the 0.25° species in the WIO).

#### 3.3.2 Composition of best models

Overall, similar predictors tended to be selected across sampling resolutions for a same sampling type (*i.e.* the same predictors were the most often selected across resolutions; Figure 8), but the predictors selected in raster-based models were not necessarily the same as for segment-based models. The model composition was variable across simulations, with few predictors being unanimously selected for a same resolution and type (few variables selected in > 70% of simulations), and the models were often mostly supported by two predictors, with a more or less variable set of supplementary predictors providing complementary information.

**Figure 8.**
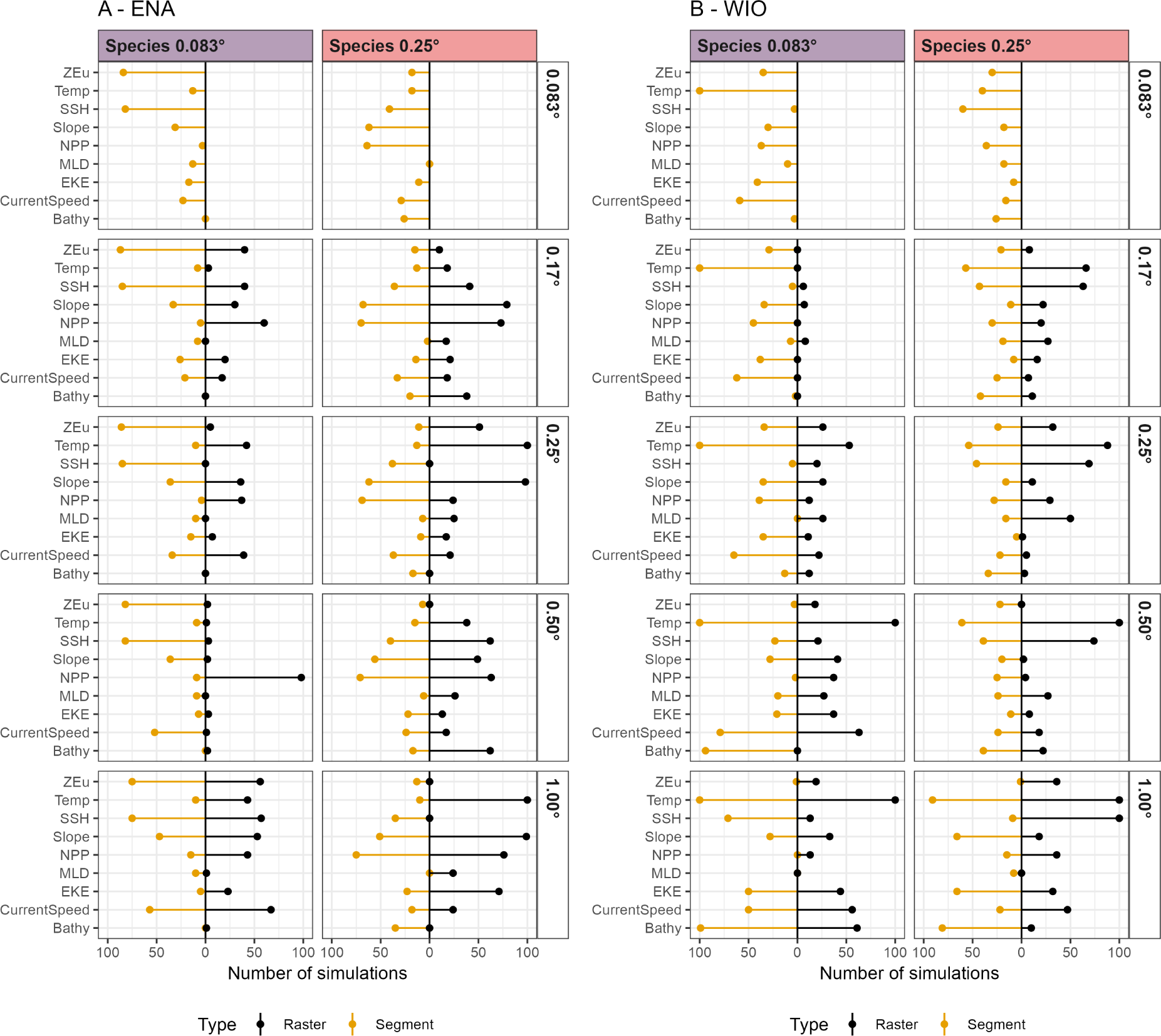
Number of simulations including each predictor in the selected best model fitted to raster-based (black) and segment-based sampling (gold), contrasting species and resolution, for (A) the ENA and (B) the WIO region. ENA: Eastern North Atlantic, WIO: Western Indian Ocean, ZEu: euphotic depth, Temp: sea surface temperature, SSH: sea surface height, NPP: net primary production, MLD: mixed layer depth, EKE: eddy kinetic energy, Bathy: bathymetry.

In the ENA, the 0.083° species segment-based models mostly selected the same predictors whatever the resolution (Figure 8), with ZEu and SSH were the two most often selected variables (> 70% of simulations), followed by Slope and CurrentSpeed (> 40% of simulations, except for the 0.083° resolution). Raster-based models were more variable in their composition, although NPP was selected in about 50% of simulations for all resolutions except the 0.50°, for which it was the sole predictor always selected. SSH, ZEu Temp, Slope and CurrentSpeed were also commonly selected in 0.17, 0.25 and 1.00° resolutions.

Similarly, for the 0.25° species in the ENA the segment-based models had similar compositions across resolutions (Figure 8), with Slope, NPP and SSH always being the three most selected predictors (> 70% of simulations for Slope and NPP, > 40% for SSH). A similar composition was found for raster-based models at the 0.17 and 0.50° resolutions, although for the latter, Bathy was often selected, unlike in the segment-based models. The composition of models for 0.17 and 1.00° were quite different however, with Temp and Slope always being selected, followed by ZEu (0.17°) and NPP and EKE (1.00°).

In the WIO, the segment-based models for the 0.083° species selected Temp whatever the resolution (Figure 8), with CurrentSpeed, EKE, Slope often selected as well (20–50% depending on resolutions). Bathy was also selected in most simulations for the 0.50 and 1.00° resolutions (90-100%). NPP was often selected as well (40–50%) but only for the lowest resolutions (0.083, 0.17 and 0.25°). For this species, raster-based models failed to find any informative predictors for all simulations at 0.17° resolution (which relates to the null explained deviances, Figure 7). At 0.25%, raster-based models were fairly variable across simulations, with Temp being the only selected in half the simulations, all other predictors being selected less than 25% of the time. At 0.50 and 1.00° Temp was systematically selected, but the remaining composition varied across simulations. Interestingly, Bathy was never selected at 0.50°, but was the second most selected covariate at 1.00° (> 50% of simulations).

As for the WIO 0.083° species, the segment-based models for the 0.25° species ended up with similar compositions at the five resolutions (Figure 8): Temp and SSH were the most selected predictors (40–50%) for all resolutions but the 1.00°, for which Temp, Slope, EKE and Bathy were selected in most of the models (> 50% of simulations for all four variables). Raster-based models all selected Temp and SSH as well, the number of simulations for which they were selected increasing with the resolution (up to 100% for the 1.00° resolution). The remaining composition of models were more variable across simulations, with all predictors being selected at some points.

#### 3.3.3 Relationships to environmental variables

Surprisingly, the relationships between the number of individuals (response variable) and the environmental conditions varied substantially across simulations in many cases. Yet, the shapes of the relationships were consistent across sampling resolutions and types for a same simulation.

The ENA 0.083° species displayed the most consistent relationships to environmental conditions across resolutions and types, with the least variable shapes across simulations (Supplementary Information S1 Figure 6). The selected best models all identified the same relationships to Temp, SSH, NPP, MLD and ZEu whatever the resolution and type. Although the shapes were broadly consistent for CurrentSpeed and EKE (except for raster-based 0.25 and 0.50° resolutions), the curves displayed more variability in their edges (where the relationships were supported by a lower amount of data). Relationships to Slope were the most variable of all environmental conditions, with no overall pattern emerging from the simulations. The ENA 0.083° species was defined from Temp, ZEu, SSH and NPP (Figure 3): all simulations selecting them found similar relationships to these four predictors, and their shapes were consistent with the true relationships. The modes of Temp and ZEu were successfully identified, but those of SSH and NPP were identified at lower values than the actual ones.

The ENA 0.25° species selected best models yielded a bit more variability across simulations, but the shapes remained consistent across resolutions and types (Supplementary Information S1 Figure 7). CurrentSpeed, MLD and EKE had the largest variability in identified relationships. This species was defined from Bathy, Temp, NPP and Slope. The relationship to Temp was successfully identified for all simulations selecting it, both in terms of shape and mode location. Although Bathy was not often selected by raster-based models, the overall shape was correctly identified when it was, despite a mode at higher depths than the actual value (−1000 m instead of −800 m). Two broad shapes of relationships to NPP were identified, one bell-shaped and one almost linear. The bell-shaped relationships were close to the true relationships, but the inflection values (of both types of relationships) were associated with lower values than what defined the species distribution (500–800 instead of about 1500). Finally, the relationships to Slope also displayed two types of shapes, one mostly linear and one alternating a minimum at low slope values and a maximum at large slope values. The second type was closer to the true relationship in its shape, but failed to identify the plateau and inflection points.

In the WIO, the inter-simulation variability in the shape of relationships was larger than for ENA species. In the case of the 0.083° species (Supplementary Information S1 Figure 8), the four variables from which it was defined (Temp, CurrentSpeed, Bathy, ZEu) were the only ones with similar shapes across simulations and resolutions (except for the raster-based 0.17° resolution which failed to select informative variables). The relationship to Temp and ZEu were successfully identified, with the inflection points correctly located, although the 0.25° raster-based models identified a bell-shaped curve instead of a plateau for Temp. The relationship to Bathy was close to the true one, but most simulations failed to identify the plateau. The raster-based models overall failed to select this variable, except at the 1.00° resolution. The relationship to CurrentSpeed was more or less correctly identified except for the segment-based 1.00° and the raster-based 0.25° models, which identified linear relationships. For this variable, the bell-shaped curve was regularly identified as a plateau because the models did not sample all the range of values used to define the species distribution (compare x axis ranges in Figure 3 and Supplementary Information S1 Figure 8)

The models for the WIO 0.25° species displayed the largest variability in the shapes of relationships identified across simulations (Supplementary Information S1 Figure 9). There was no consistent pattern in relationships for Bathy, Slope, NPP, MLD, CurrentSpeed and Zeu, with some relationships completely opposite for two different simulations at the same resolution and type (*e.g.* CurrentSpeed in segment-based models). Temp and SSH were the only two variables for which simulations all agreed upon the shape of relationship. These were two of the variables used to define the species, and the shapes were consistent with the truth, despite the left bell-shape often identified as a plateau and the inflection points identified at larger values than the actual ones. MLD and NPP were the other two variables used to define the species. Only some simulations broadly identified the overall shape of the relationship to MLD (raster-based models only), while all models failed to identify the relationship to NPP.

**Figure 9.**
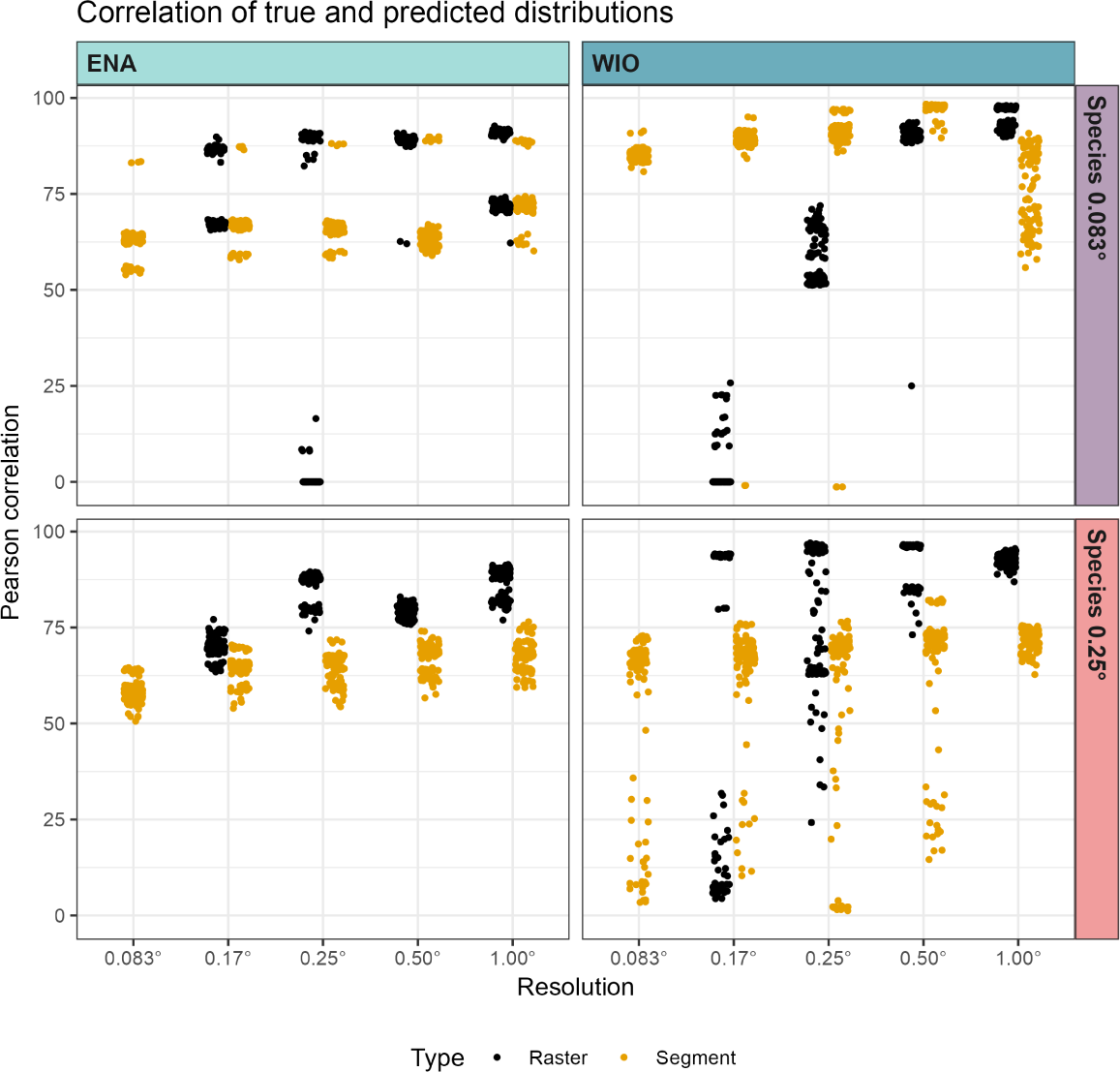
Distribution of the Pearson’s correlation coefficient between the true and predicted abundance maps. Predicted abundances were derived from best models fitted to raster-based (black) and segment-based sampling (gold), contrasting species and resolution, for the ENA and the WIO regions. ENA: Eastern North Atlantic, WIO: Western Indian Ocean.

### 3.4 Model predictions

#### 3.4.1 Quality of predictions

As the composition of best models were variable, so were the predicted distributions. In line with the more stable composition of segment-based models, the correlation of their prediction with the true distribution did not quite change across resolutions for the two ENA species (always larger than 50%), although correlations were somewhat higher for larger resolutions (Figure 9A). The corresponding raster-based models yielded better correlations in most cases, with the exception of the 0.17° resolution for the 0.083° species where some simulations fared badly due to poor quality models (null model selected as the best one). For this species, all other models whatever resolutions and types split into two bulks of correlation values, at 55–70% and at 80–90%, respectively. This splitting suggested that some simulations (*i.e.* some set of observations) were more informative than others, resulting in higher quality predictions.

For the WIO species however, the best models were less successful in replicating the true distributions (Figure 9A). In the case of the 0.083° species, segment-based models had good correlation values for the 0.083–0.50° resolutions (80– 96%), but the correlations drop at 1.00°, with a large variability. The raster-based models yielded very different results. In line with the very low to null deviances of selected models, the prediction poorly related to the true distribution for the 0.17° resolution: most simulations yielded null correlations (null model selected as best one), and the few others which selected covariates in the best models performed very poorly (< 25%). The correlations were variable but higher for the 0.25° resolution, and high for the 0.50 and 1.00°. For the latter resolution, the raster-based models fared clearly better than the segment-based ones.

As models struggled to identify consistent relationships to environmental conditions, both segment-based and raster-based models struggled to replicate the true distribution of the WIO 0.25° species (Figure 9A). No segment-based models exceeded the 75% of correlation, and a non-negligible portion of simulations entirely failed to reproduce the distribution (correlations lower than 50%), for all resolutions but the 1.00° one. These low correlations were obtained by simulations for which models yielded aberrant predictions in some cells of the map (*i.e.* predicted extreme and unrealistic density; not shown). Raster-based models at the 1.00° resolution provided the best results (correlation larger than 80%), alongside a handful of simulations at the 0.17, 0.25 and 0.50° resolutions. For these, the variability in correlation values across simulations was the widest, denoting an overall poor quality of the models. Again, as for the 0.083° species in the ENA, this pattern indicated that some simulations were more informative than others for the model to reconstruct the species distribution.

#### 3.4.2 Predicted spatial distributions

To get an overview of the predicted spatial patterns, we averaged together the predictions obtained for the 100 simulations for each region, species, type and resolution. It must be reminded that an extensive variability in predicted pattern across simulations might be masked by the averaging procedure (in particular, the aberrant and extreme pixels in individual predictions are buffered and not seen). For the particular cases of the 0.083° species in the ENA sampled at the 0.25° raster-based resolution, and of the 0.083° species in the WIO sampled at the 0.17° raster-based resolution, the maps displayed in Figure 10 included the 52 and 82 simulations (respectively) where the null model was selected as best one (predicting homogeneous maps of null density). Hence, we do not discuss these results in the following.

**Figure 10.**
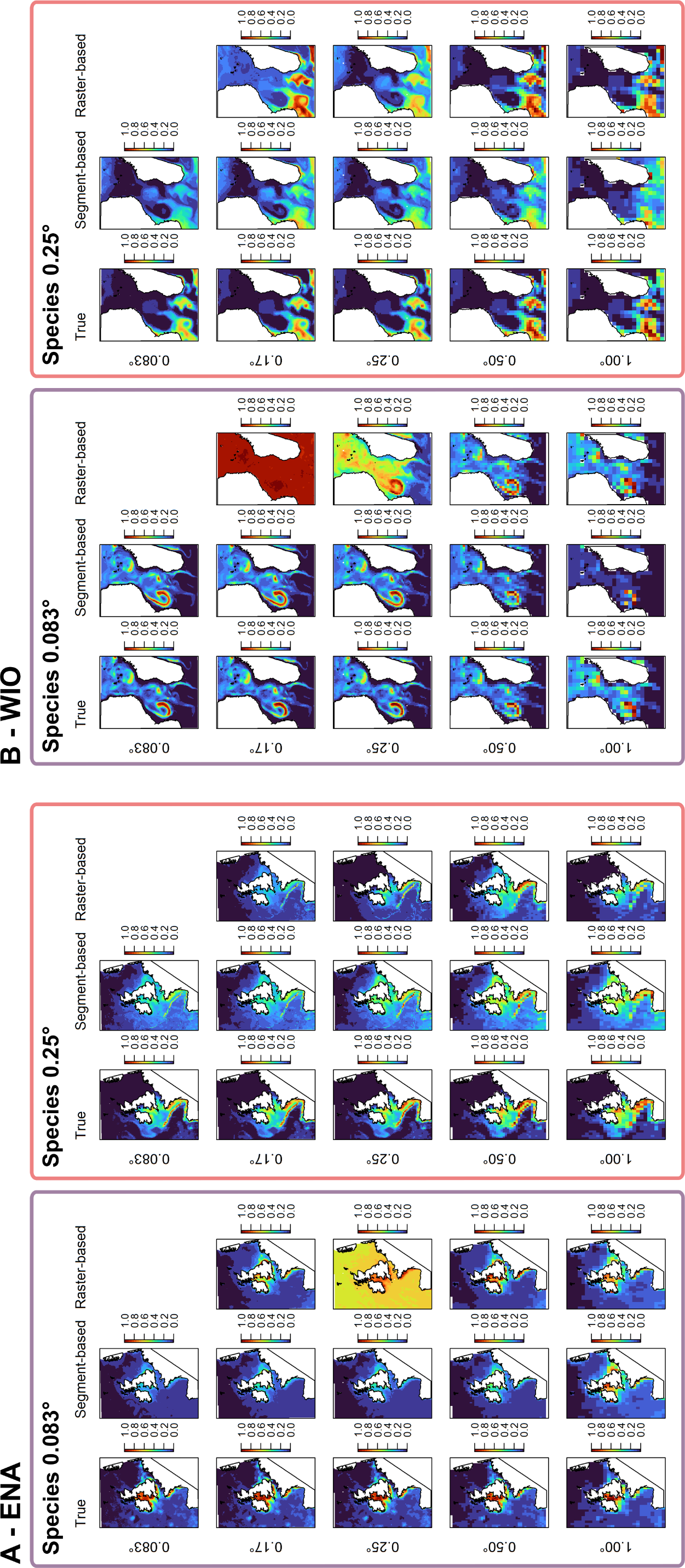
Species true and predicted distributions averaged over the 100 simulations for each sampling type and sampling resolution (as relative density), for the 0.083° species (left) and the 0.025° species (right) in the ENA (A) and WIO (B). ENA: Eastern North Atlantic, WIO: Western Indian Ocean. The true distributions correspond to the true density of individuals per cell averaged over the 100 simulations.

In the ENA (Figure 10A), the true distribution pattern for the 0.083° species was mostly found for all resolutions and types (except for the above-mentioned raster-based 0.25°), despite too high densities often predicted along the English coast of southern North Sea. The models also underestimated the density in the central Irish Sea (especially for segment-based models), but the distribution patterns were well predicted in the Bay of Biscay shelf. Overall however, the models did not captured the small-scale low-density patterns occurring in offshore waters. For the 0.25° species (Figure 10A), the overall preference for the slope was correctly predicted whatever resolutions and types, but models were less successful in predicting the true distribution pattern over the shelf: medium densities were predicted in the southern North Sea and Eastern Channel while true occurrences there were close to null. Distribution pattern in the Irish Sea and Bay of Biscay were well predicted, but not in the Iberian shelf. Overall, models predicted too high densities in the oceanic areas (*i.e.*, beyond shelf edge) but the patch of higher density over the seamount off Galicia was predicted for all resolutions and types. In average, the 0.25 and 1.00° resolutions provided the best predictions for both raster and segment-based types.

In the WIO (Figure 10B), the segment-based models for the 0.083° species all found similar patterns, although they diluted as the resolution enlarged, resulting in the 1.00° resolution oversimplifying the pattern. For the other segment-based resolutions, the pattern was in average correctly identified. The large-scale latitudinal gradient as well as the large densities associated with oceanic features were well retrieved (eddies and filaments), despite overestimated densities along the western edge of the Mozambique Channel eddy for the 0.083, 0.17 and 0.25° resolutions. The raster-based models at 0.25 resolutions failed to find a consistent pattern across the simulations. At the 0.50° resolution, raster-based models provided similar results as segment-based models, while at the 1.00° resolution, they predicted spatial patterns strongly consistent with the true distribution, unlike segment-based models.

Although most models found the distribution patterns in the south of the WIO region for the 0.25° species (Figure 10B), all models also predicted medium densities in the northern and central parts of the study area, as well as in between features hosting the largest densities, despite very low true densities there. Segment- and raster-based models predicted similar patterns for each resolutions, except at 1.00° resolution where the segment-based predictions were less accurate than the raster-based ones. Overall however, these averaged patterns might be largely influenced by the models whose performances were the lowest (Figure 9).

## 4 Discussion

In the present work, we implemented an unprecedented simulation procedure to inspect the effects of two overlooked and misunderstood aspects of SDM: the choice of environmental condition resolution and the choice of sampling type (segment *vs* raster-based) used to fit the models. The performances of SDMs were inspected through various means, from statistical metrics to prediction quality. This study is, to our knowledge, the first to quantify the effect of sampling type on model performances, and our simulation approach allowed us to demonstrate for the first time the variability of the modelling process in curve complexity and model compositions. Thanks to the use of two different study areas and two different virtual species, we disentangled the effects of species selectivity, environmental heterogeneity, sampling types, sampling resolutions and survey stochasticity on the ability of a SDM to reliably estimate and reproduce the true species distribution.

### 4.1 Methodological limitations

This study was based on simulations built to match as much as possible real-life cases of species distribution modelling exercises. Yet, as all simulations, they came with several technical choices and limitations.

First, we simulated a survey using a strip-transect protocol, hence assuming perfect detection of animals (Buckland et al., 2015). This is never the case in real life, where many factors alter detection probability. The detection probability of animals is directly dependent on natural factors affecting the observer ability to see items present within the surveyed area. Such factors are not specific to surveys carried out in the marine domain, and can range from sunlight intensity, sun angle, to cloud cover (for both terrestrial and marine surveys) or sea state (for marine surveys). In addition to natural factors pertaining to the environment in which the observations are carried out, the detection probability is also conditioned by the behaviour of the animals actually sighted (*i.e.* animals are not always available for detection). In terrestrial domain, hiding or burying behaviours hinder the detectability of animals by observers, while in the marine realm, it is the ability of animals to dive that do so. In real-life cases, these parameters are routinely taken into account in the analysis through correction factors. For example, the abundance estimated from aerial surveys for diving marine species can be corrected by the time they spend at the sea surface. Such process would reduce the actual number of sightings, potentially not homogeneously in space and time, and add some noise in the dataset that could potentially affect the ability of a modelling procedure to find relevant models.

We simulated a very large-scale survey but ignored the logistical constraints associated with such. In particular, we ignored the need for refuelling, which would make the most remote parts of the study areas practically impossible to survey. Furthermore, we simulated a single-day survey, which is completely unrealistic given the size of the study areas. In real-life cases, surveys of that extent would necessitate a non-negligible number of aircraft and crews, and would take up to several months to be completed.

This strong choice was made with the aim of simulating a simultaneous survey, with all segments observed at the same time. Doing this, we could ignore the movements of the target species. Indeed, large sized marine species, which are the main target of aerial and boat-based surveys in the marine domain, are extensively mobile and can move within very large home ranges during relatively short periods of time (shorter time period than survey durations). The effect this discrepancy between the scale of the individual home ranges and that of the survey can have on the output of SDMs is still misunderstood, so we chose to ignore the movement of animals in our simulation and used an IPPP, *i.e.* simulating individuals with static positions. In addition to ignoring animal movements, we also ignored the aggregative behaviour that most marine species exhibit by simulating single individual locations. However, we anticipate this choice did not have much of an impact on our results since the number of individuals sighted was summarized per segment, and this final number of individuals would be similar if they were sighted together or separately.

The abundance and prevalence of species are known to have a considerable impact on the ability to build relevant and reliable SDMs (Hernandez et al., 2006; Virgili et al., 2018). The distribution of species with low abundance (*i.e.* “rare”) is notoriously difficult to model, due to the very low proportions of sampling units with presences (the zero-inflation problem). Yet, the prevalence of a rare species is what really determines if a statistical model is able to identify the underlying drivers and patterns: a rare species with low prevalence and marked habitat preferences (high selectivity) will be more easily modelled and predicted (Virgili et al., 2018, 2019), in particular when occurring in heterogeneous environment (Connor et al., 2017), than a rare but widespread species. Here, we simulated four species with the same abundance and prevalence (moderate) so that the results were directly comparable across species. Our results would probably remain valid for more abundant species, but would probably be more variable if we were to simulate a truly rare species. The effect of the species prevalence however should be largely dependent on the characteristics of the study area, and in particular upon its environmental heterogeneity.

### 4.2 The effect of sampling type

The single variable modelling provided clear evidence that, for a single simulated dataset, using raster-based sampling yielded higher deviances, but this statistical performance improvement came at the cost of simplifying the identified relationships and decreasing predictive performance (lower RMSE). This held true for all the considered environmental conditions, species and regions and whatever the sampling resolution used. In addition, deviances explained by each environmental condition separately were highly variable depending on simulations when using raster-based sampling, while explained deviances remained consistent across simulations for segment-based sampling.

The output of the model selection procedure proved to be surprisingly variable across simulations for both sampling types. Although all simulations were based on the same occurrence maps, the results sometimes differed greatly across datasets, highlighting a certain level of stochasticity in the modelling process, originating from the raw data. As observed with single variable models, raster-based final models most often ended up in simpler (linear) relationships to environmental conditions, failing to identify complex features of the original species-condition relationship. Segment-based models were better at retrieving this original relationship, although the inflection points were often misidentified. This pattern was more striking at larger resolutions, which may be linked to the lower number of sampling units available to fit raster-based models at these resolutions compared to segment-based ones: the amount of data directly influences the spline parameter identification in the GAM (Wood, 2006). Additionally, we can observe a difference in the stochasticity of curve shapes between the two oceanographic regions, the WIO models having more variable curve shapes across simulations than the ENA ones.

Interestingly, most of the deviance in final models was explained by one or two predictors (not shown here, but models with more variables did not achieve better correlation between true and predicted distributions). Yet, the variables selected in best models were not necessarily the ones used to simulate the species distribution. This is due to the GAMs being correlative models (Wood, 2006): the best models select predictors best reproducing the spatial distribution of the data, not that better explain it. The discarding of models including pairs of too correlated predictors when selecting best models might also be involved in this pattern. In some cases, one of the conditions used to simulate the species have been discarded because of their correlation with another one. This discarding of correlated variables may also drive the distribution of explained deviances into “bulks” of similar values (also true for correlation between prediction and true spatial distribution): one condition is selected against another in some simulations, while the selection pattern reverses for other ones, resulting in different model performances.

Overall, our simulations clearly emphasized and confirmed the necessity not to over-interpret the environmental conditions selected as best predictors in final species distribution models.

### 4.3 The effect of the environmental heterogeneity

Surprisingly, and unlike what has been previously observed in the literature (Guisan et al., 2007; Connor et al., 2017), our simulation study highlighted no clear adequation between the performance of sampling resolution and the true resolution of environmental drivers: the spatial patterns of distribution predicted at the resolution closest to the one used to simulate the species did not stand out compared to models using either larger or smaller resolutions. Our simulated species and study areas were relatively resistant to the coarsening of resolution. Yet, as hinted by the larger stochasticity in curve shapes and prediction quality for the WIO region, the model sensitivity to the resolution of environmental conditions seems to depend on the oceanographic processes underlying species distributions.

In the ENA, the oceanographic processes used to simulate the species at both 0.083° and 0.25° resolutions (tidal and slope-associated structures) remained identifiable at larger resolution. This resulted in the models being able to identify the distribution patterns whatever the resolution, and the inconsistencies between occurrence and predicted maps to mostly derive from the model abilities. On the contrary, in the WIO, the fine-scale processes selected by the species (edges of meso-scale structures) were diluted when the resolution was increased, especially beyond 0.50°. This has different implications depending on the sampling type considered. For segment-based sampling, the fine-scale processes disappeared from the environmental conditions maps as the resolution increase, so that the models lose informative signal in the predictors and failed to identify reliable drivers (coarsening the resolution reduce the informative level of predictors). For raster-based sampling however, the models failed to identify environmental conditions successfully reproducing the spatial pattern because this very same spatial pattern in species distribution was altered by the rasterisation process. In this case, the low performances of models essentially lie in the low informative levels of the input data (response variable).

These results thus shed a new light on the effect of sampling resolution on SDM performance. We knew this performance depends on the species prevalence, habitat selectivity (specialist vs generalist) and the environment heterogeneity (Hernandez et al., 2006; Gottschalk et al., 2011; Lauzeral et al., 2013; Connor et al., 2017), with the alterative effect of coarsening the resolution reduced for specialist species in heterogeneous environment. Yet, here, by controlling the species prevalence and abundance, we demonstrate that the effect of coarsening resolution is mostly linked to the resistance of the environmental features selected by the species to this coarsening, rather than to the intrinsic heterogeneity of the complete landscape. As a result, SDM would be less sensitive to the choice of sampling resolution when used for species targeting environmental features that are well-preserved when changing the spatial resolution. It could be advisable, then, to test for the preservation of the observed structures in species spatial patterns before choosing the resolution at which the model will be built.

Yet, the higher robustness of segment-based models on coarsening the resolution indicated that the choice of sampling type (segment- or raster-based) had more effects on the final SDM performance than the actual resolution used in the model, due to the risk of losing information when rasterising observation data (*i.e.* plummeting the information contained in the response variable). And, more importantly and surprisingly, our results demonstrated without ambiguity the potentially large effect of the stochasticity inherent to the raw dataset in the final performance of the models. Indeed, the across-simulation variability was often larger than that associated with the resolution or type chosen to model the species. This pattern might explain why, in some real-life cases, all approaches fail to fit successfully an adequate and relevant model. We therefore urge to caution when setting up modelling studies and interpreting their results.

### 4.4 Take-home message

Our simulation study provided clear evidence of stochasticity in the modelling process, thereby urging modellers to caution in fitting models and interpreting resulting outcomes. We also evidenced that classical statistical performance metrics (explained deviances, RMSE. . .) are not good correlates of predictive quality (for spatial pattern). Despite raster-based modelling being faster in computation (thanks to the lower amount of data points), segment-based models seemed to be more robust to changes in predictor resolution, and are to be preferred as this approach also avoids the potential loss of information that could occur when rasterising at scales larger than that of biological significance. However, this pattern is strongly dependent on how much the species distribution and/or environmental conditions are impacted by changes in scale. We therefore advise checking for the resistance of spatial patterns to changes of resolution, both for response and predictor variables, before any analysis. To test for this, it may be considered to quantify the heterogeneity of both the environment and the species distribution, at different resolutions, with tools rooted in landscape ecology. Furthermore, if one decides to opt for raster-based analysis, we strongly recommend carefully checking that the spatial patterns observed with the segmented data are still clearly identifiable after the rasterisation process. Above and foremost, the final choice of sampling type and resolution must depend on the question at hand. This choice must be informed carefully, so that the scale is, as much as feasible, adequately tuned between the observation process and the environmental predictors.

## 5 Supplementary information and data availability

Supplementary Information, data and R codes used to complete the analyses presented in this paper are accessible in a dedicated repository on Zenodo, accessible at https://doi.org/10.5281/zenodo.7544441.

## References

Baddeley A, Rubak E, Turner R, 2015 Spatial Point Patterns: Methodology and Applications with R (London: Chapman and Hall/CRC Press) URL http://www.crcpress.com/Spatial-Point-Patterns-Methodology-and-Applications-with-R/Baddeley-Rubak-Turner/9781482210200/

Buckland S, Borchers D, Marques T, Fewster R, 2023 “Wildlife population assessment: Changing priorities driven by technological advances” Journal of Statistical Theory and Practice 17 20

Buckland S, Rexstad E, Marques T, Oedekoven C, 2015 Distance sampling: methods and applications (Springer)

Caballero A, Ferrer L, Rubio A, Charria G, Taylor B H, Grima N, 2014 “Monitoring of a quasi-stationary eddy in the Bay of Biscay by means of satellite, in situ and model results” Deep-Sea Research II 106 23–37

Connor T, Hull V, Viña A, Shortridge A, Tang Y, Zhang J, Wang F, Liu J, 2017 “Effects of grain size and niche breadth on species distribution modeling” Ecography 41 1270–1282

de Ruijter W P, van Aken H M, Beier E J, Lutjeharms J R, Matano R P, Schouten M W, 2004 “Eddies and dipoles around South Madagascar: formation, pathways and large-scale impact” Deep Sea Research Part I: Oceanographic Research Papers 51 383–400

Elith J, Leathwick J R, 2009 “Species Distribution Models: Ecological Explanation and Prediction Across Space and Time” Annual Review of Ecology, Evolution, and Systematics 40 677–697

Fernandez M, Yesson C, Gannier A, Miller P I, Azevedo J M, 2017 “The importance of temporal resolution for niche modelling in dynamic marine environments” Journal of Biogeography 44 2816–2827

Franklin J, 2010 Mapping species distributions: spatial inference and prediction (Cambridge University Press)

Franklin J, 2013 “Species distribution models in conservation biogeography: developments and challenges” Diversity and distributions 19 1217–1223

Gottschalk T K, Aue B, Hotes S, Ekschmitt K, 2011 “Influence of grain size on species–habitat models” Ecological Modelling 222 3403–3412

Guillera-Arroita G, Lahoz-Monfort J J, Elith J, Gordon A, Kujala H, Lentini P E, McCarthy M A, Tingley R, Wintle B A, 2015 “Is my species distribution model fit for purpose? Matching data and models to applications” Global Ecology and Biogeography 24 276–292

Guisan A, Graham C H, Elith J, Huettmann F, Group N S D M, 2007 “Sensitivity of predictive species distribution models to change in grain size” Diversity and distributions 13 332–340

Guisan A, Tingley R, Baumgartner J B, Naujokaitis-Lewis I, Sutcliffe P R, Tulloch A I, Regan T J, Brotons L, McDonald-Madden E, Mantyka-Pringle C, et al., 2013 “Predicting species distributions for conservation decisions” Ecology Letters 16 1424–1435

Harrell Jr F E, with contributions from Charles Dupont, many others., 2020 Hmisc: Harrell Miscellaneous r package version 4.4-2 URL https://CRAN.R-project.org/package=Hmisc

Hernandez P A, Graham C H, Master L L, Albert D L, 2006 “The effect of sample size and species characteristics on performance of different species distribution modeling methods” Ecography 29 773–785

Hijmans R J, Etten J v, Mattiuzzi M, Sumner M, Greenberg J A, Lamigueiro O P, Bevan A, Racine E B, Shortridge A, 2014 “raster: Geographic data analysis and modeling” URL http://cran.r-project.org/web/packages/raster/index.html

Koutsikopoulos C, Le Cann B, 1996 “Physical processes and hydrological structure related to the Bay of Biscay anchovy” Scientia Marina 60 9–19

Lambert C, Authier M, Blanchard A, Dorémus G, Laran S, Van Canneyt O, Spitz J, 2022 “Delayed response to environmental conditions and infra-seasonal changes in the spatial distribution of short-beaked common dolphin” Royal Society Open Science

Lambert C, Authier M, Dorémus G, Gilles A, Hammond P, Laran S, Ricart A, Ridoux V, Scheidat M, Spitz J, et al., 2019 “The effect of a multi-target protocol on cetacean detection and abundance estimation in aerial surveys” Royal Society open science 6 190296

Lambert C, Mannocci L, Lehodey P, Ridoux V, 2014 “Predicting cetacean habitats from their energetic needs and the distribution of their prey in two contrasted tropical regions” PLoS One 9 e105958

Lauzeral C, Grenouillet G, Brosse S, 2013 “Spatial range shape drives the grain size effects in species distribution models” Ecography 36 778–787

Leroy B, Meynard C N, Bellard C, Courchamp F, 2015 “virtualspecies, an r package to generate virtual species distributions” Ecography 6

Longhurst A R, 2007 Ecological geography of the sea 2nd edition (Academic Press)

Mannocci L, Boustany A M, Roberts J J, Palacios D M, Dunn D C, Halpin P N, Viehman S, Moxley J, Cleary J, Bailey H, et al., 2017 “Temporal resolutions in species distribution models of highly mobile marine animals: Recommendations for ecologists and managers” Diversity and Distributions 23 1098–1109

Mannocci L, Catalogna M, Dorémus G, Laran S, Lehodey P, Massart W, Monestiez P, Van Canneyt O, Watremez P, Ridoux V, 2014a “Predicting cetacean and seabird habitats across a productivity gradient in the South Pacific gyre” Progress in Oceanography 120 383–398

Mannocci L, Laran S, Monestiez P, Dorémus G, Van Canneyt O, Watremez P, Ridoux V, 2014b “Predicting top predator habitats in the Southwest Indian Ocean” Ecography 37 261–278

Manzoor S A, Griffiths G, Lukac M, 2018 “Species distribution model transferability and model grain size–finer may not always be better” Scientific reports 8 1–9

Marshall C, Glegg G, Howell K, 2014 “Species distribution modelling to support marine conservation planning: The next steps” Marine Policy 45 330–332

Marshall L, 2021 dssd: Distance Sampling Survey Design r package version 0.3.1 URL https://CRAN.R-project.org/package=dssd

Moudry V, Keil P, Cord A F, Gábor L, Lecours V, Zarzo-Arias A, Barták V, Malavasi M, Rocchini D, Torresani M, et al., 2023 “Scale mismatches between predictor and response variables in species distribution modelling: A review of practices for appropriate grain selection” Progress in Physical Geography: Earth and Environment 03091333231156362

Pingree R D, Le Cann B, 1992 “Three anticyclonic Slope Water Oceanic eDDIES (SWODDIES) in the southern Bay of Biscay in 1990” Deep Sea Research 39 1147–1175

R Core Team, 2021 R: A Language and Environment for Statistical Computing R Foundation for Statistical Computing Vienna, Austria URL https://www.R-project.org/

Scales K L, Hazen E L, Jacox M G, Edwards C A, Boustany A M, Oliver M J, Bograd S J, 2016 “Scale of inference: on the sensitivity of habitat models for wide-ranging marine predators to the resolution of environmental data” Ecography

Schouten M W, de Ruijter W P, Van Leeuwen P J, Ridderinkhof H, 2003 “Eddies and variability in the Mozambique Channel” Deep Sea Research Part II: Topical Studies in Oceanography 50 1987–2003

Spiess A N, 2018 qpcR: Modelling and Analysis of Real-Time PCR Data r package version 1.4-1 URL https://CRAN.R-project.org/package=qpcR

Virgili A, Authier M, Monestiez P, Ridoux V, 2018 “How many sightings to model rare marine species distributions” PLoS ONE 13 e0193231

Virgili A, Racine M, Authier M, Monestiez P, Ridoux V, 2017 “Comparison of habitat models for scarcely detected species” Ecological Modellling 346 88–98

Virgili A. and Authier M, Boisseau O, Cañadas A, Claridge D, Cole T, COrkeron P, Dorémus G, David L, Di-Méglio N, Dunn C, Dunn T, Garcia-Baron I, Laran S, Lauriano G, Lewis M, Louzao M, Mannocci L, Matinez-Cedeira J, Palka D, Panigada S, Pettex E, Roberts J, Ruiz L, Saavedra C, Santos M, Van Canneyt O, Vazquez Bonales J, Monestiez P, Ridoux V, 2019 “Combining multiple visual surveys to model the habitat of deep-diving cetaceans at the basin scale” Global Ecology and Biogeography 00 1–15

Wood S, 2006 Generalized Additive Models: An Introduction with R 1st edition (Boca Raton, FL: Chapman and Hall/CRC)

Wood S N, 2011 “Fast stable restricted maximum likelihood and marginal likelihood estimation of semiparametric generalized linear models” Journal of the Royal Statistical Society: Series B (Statistical Methodology) 73 3–36

